# Machine Learning for Toxicity Prediction in Low-Sample Molecular Classes

**DOI:** 10.64898/2026.09.16.751426

**Authors:** Carlos Barajas, Laura Dunphy, Luke Mullany, Olivia Tiburzi, Evan Lloyd

## Abstract

Deep learning models such as Chemprop have advanced quantitative molecular property prediction, but their reliance on large training sets limits use in data-scarce domains. We propose a framework that fine-tunes a general baseline model trained on publicly available data on small, class-specific datasets. The resulting models retain the baseline’s generalization ability while gaining class-specific accuracy and produce probabilistic outputs that capture uncertainty in the training data. We demonstrate the approach on three toxicity classes defined by a common core structure, target, or mode of action: (i) organophosphates, (ii) androgen receptor antagonists, and (iii) estrogen receptor *β* antagonists. Each fine-tuned model outperforms classical machine-learning methods and the EPA TEST tool. The probabilistic nature of the predictions enables prioritization of compounds for experimental validation and seamless integration with data streams of varying quality, supporting iterative decision-making in chemical safety and drug discovery.

## 1 Introduction

Accurate prediction of quantitative molecular properties from chemical structure underpins high-impact applications such as early-stage pharmaceutical lead optimization, generative AI-driven exploration of chemical space, and rapid assessment of next generation pesticides.^1–5^ In particular, toxicity-prediction models aid in screening pharmaceutical drug designs and serve as an ethical alternative to experimental methods that commonly determine the median Lethal Dose (LD_50_) of the molecule on a population of animals.^6,7^

In practical applications, scientists are often interested in developing prediction models for a specific molecular class of interest rather than using generalized models.^8^ In this work, we consider target-based and structure-based molecular classes of interest. A target-based class is defined by a shared biological target or mode of action, such as those tailored to a particular receptor or target family. Comparatively, a structure-based class of interest is defined by shared core structure(s) that define a chemical family. However, a key challenge in developing such class-specific predictive models is the limited availability of relevant training data.^9^ This limitation is especially pronounced in novel drug-discovery efforts, where the chemical diversity of candidate molecules is rapidly expanding due to the rise of generative AI approaches.^5,10,11^ Moreover, uncertainty stems not only from noisy measurements due to biological variance but also from the fusion of datasets collected under different experimental conditions; alignment of such heterogeneous sources with simple mappings (e.g., linear regression) adds error that propagates as epistemic uncertainty. Consequently, there is a clear need for class-specific models that retain high accuracy on the molecular class of interest, explicitly handle this uncertainty, and enable rapid, cost-effective active-learning screening workflows.^12,13^

Quantitative Structure-Activity Relationship (QSAR) models were originally pioneered by building statistical and machine-learning models on fixed molecular descriptors and fingerprints, then testing various regressors such as random forests or support-vector machines.^14,15^ The deep-learning boom later shifted the paradigm toward neural networks that learn data-driven representations (unique “fingerprints”) directly from Simplified Molecular Input Line Entry System (SMILES) or molecular graphs and couple them to a downstream regressor, improving predictive fidelity but increasing data-requirement demands.^16,17^ Consequently, sparse molecular classes with limited samples render deep models less applicable, motivating the use of zero-shot, few-shot, and transfer-learning strategies to leverage knowledge from related tasks.^9,12^ Meanwhile, uncertainty quantification and active-learning loops have been introduced to handle noisy measurements, heterogeneous metadata, and to prioritize informative compounds for rapid screening,^13,18,19^ underscoring the need for QSAR pipelines for molecules sharing a common biological target, mode of action, or structural feature that are both accurate and robust to uncertain data.

To address the limited size of class-specific data sets and the uncertainty introduced by merging heterogeneous toxicology data, we propose a unified three-step framework that goes beyond the standard pre-train / fine-tune pipeline.

1. **Global Pre-Training.** A deep neural network is trained on the full public compound library to learn general chemical representations.
2. **Class-Focused Fine-Tuning with Similarity-Curated Data.** For a given user-defined molecular class (e.g., target-based, structure-based) the baseline model is fine-tuned not only on the scarce class examples but also on a set of structurally similar compounds drawn from the global pool. This similarity-guided augmentation supplies relevant chemical context while keeping the model focused on the target biology.
3. **Probabilistic Head via Gaussian-Process Regression (GPR).** A GPR layer is added and jointly fine-tuned, providing predictive uncertainties and a natural acquisition function for active-learning loops. The probabilistic head explicitly captures error from noisy measurements and from the mapping of heterogeneous metadata.

The novelty of our approach lies in (a) the use of similarity-selected auxiliary data during the fine-tuning stage, (b) the integration of a GPR uncertainty head on top of a deep-learning backbone, and (c) the seamless incorporation of the resulting model into prospective molecular screening workflows that require rapid, cost-effective decision making and active learning.^12, 13^

We evaluate the three-step framework on three low-resource toxicity classes of interest, each with fewer than *≈*250 labeled compounds: (i) organophosphates (OPs), (ii) androgen receptor (AR) antagonists, and (iii) estrogen receptor *β* (ER*β*) antagonists. Baseline performance for each class is measured with the United States Environmental Protection Agency (U.S. EPA) gold-standard toxicity model, Toxicity Estimation Software Tool (TEST).^20^ The EPA model consistently outperforms any fixed-fingerprint representation combined with a downstream regressor across a wide hyper-parameter grid, and it also exceeds deep-learning models trained solely on the limited class-specific data. Adding the similarity-guided fine-tuning and the Gaussian-process regression head further improves prediction accuracy for all three classes and provides calibrated uncertainty estimates that can be used in active-learning-driven screening.^12,13^

## 2 Methods

In this section we present the methods to create uncertainty-aware (Figure 1-a) class-specific machine learning QSAR models for user-defined molecular classes that can aid in experimental prioritization (Figure 1-b and SI section *Bayesian Optimization for Molecular Screening* for more details). First, we introduce the dataset used for model building on this effort. Then, we review our model training framework (Figure 1-c). Finally, we discuss the methods used to benchmark performance of these toxicity prediction models.

**Figure 1:**
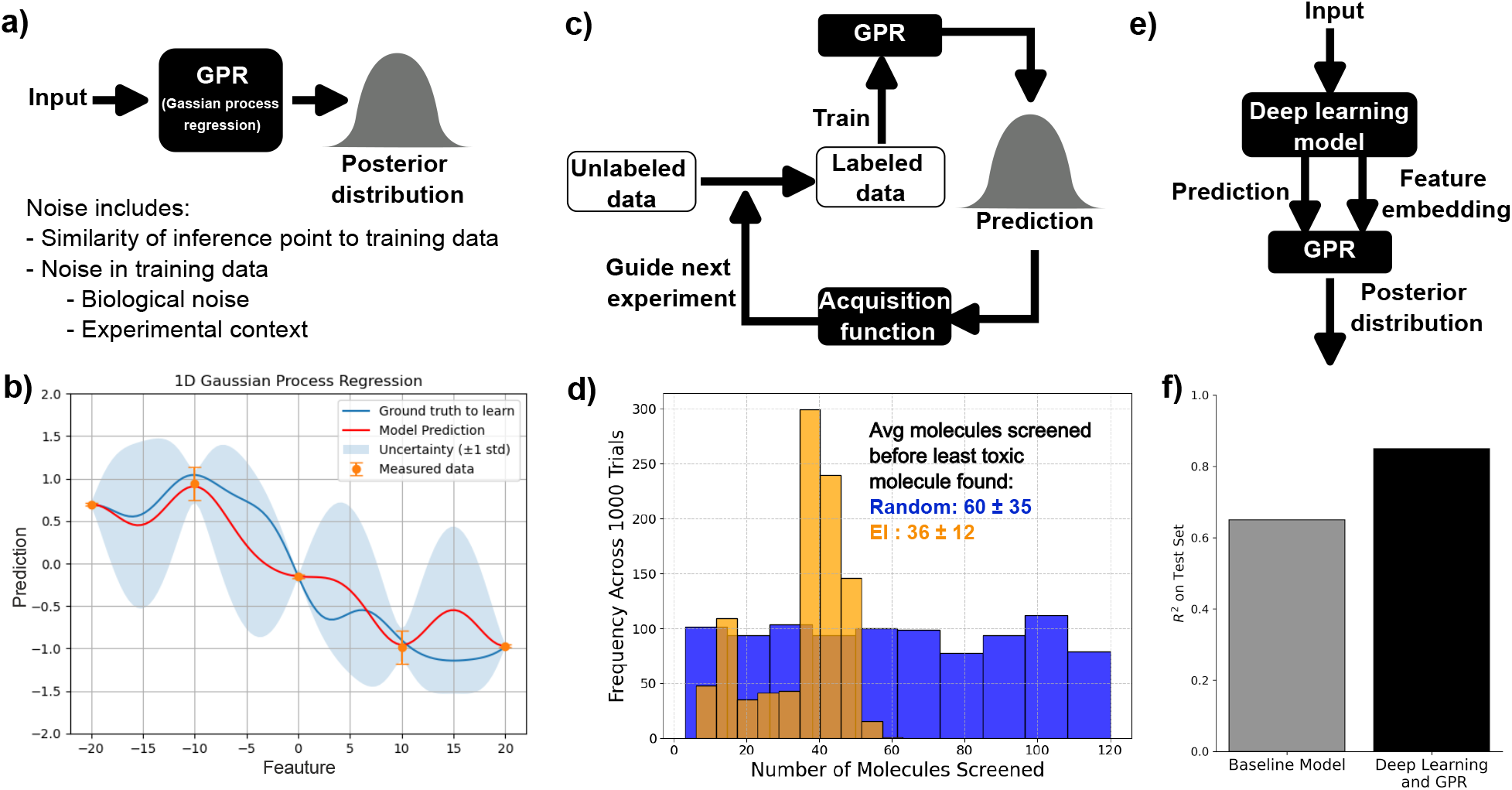
Gaussian Process Regression (GPR) for Probabilistic Regression and Active Learning. (**a**) GPR models provide probabilistic predictions that capture both the noise in the training data and the distance of inference points from that data. (**b**) One-dimensional toy illustration: high uncertainty is observed for points far from the training set, whereas regions with dense training data show low uncertainty. (**c**) By quantifying predictive uncertainty, GPR enables rational prioritization of experiments and guides the selection of the most informative measurements. (**d**) Using GPR to search for a molecule with the lowest toxicity from an unlabeled pool of candidate molecules, is 1.7 times faster than a random-sampling strategy (see SI section *Bayesian Optimization for Molecular Screening* for more details). (**e**) When coupled with deep-learning models, GPR delivers high-fidelity, classspecific predictions for stucure-defined or target-defined molecular classes. (**f** ) This work demonstrates that a synergistic model, combining a deep-learning estimator with a GPR layer, outperforms baseline approaches while retaining all the advantages of Gaussian-process uncertainty quantification.

### 2.1 Datasets

In this section, we describe the general toxicity dataset used in this work, as well as the one structure-defined and two target-defined molecular classes for which we build class-specific models.

#### 2.1.1 Canary Dataset

The Canary tool is a computational toxicology tool capable of predicting a large number of toxicology endpoints as well as physical properties, developed by Pacific Northwest national Laboratory (PNNL) for the Department of Homeland Security (DHS) Science and Technology Directorate (S&T). Each assessment includes a similarity search against the Canary database, a comprehensive collection of *>* 900k compounds with experimentally measured toxicological endpoints gathered from various public sources. In this work we refer to a frozen snapshot of that collection as the “Canary dataset”.

The dataset is in tabular form and column descriptions are provided in Table 1. We solely concentrated on oral LD_50_ data as a proof-of-concept for our approach, though the datatset includes many toxicity endpoints and routes of administration (ROAs). Mouse was taken as the primary species, and rat data were added only for SMILES that did not appear in the mouse set. After filtering the Canary dataset by administration route and animal test subjects, the resulting dataset contains approximately 28,000 compounds.

**Table 1:**
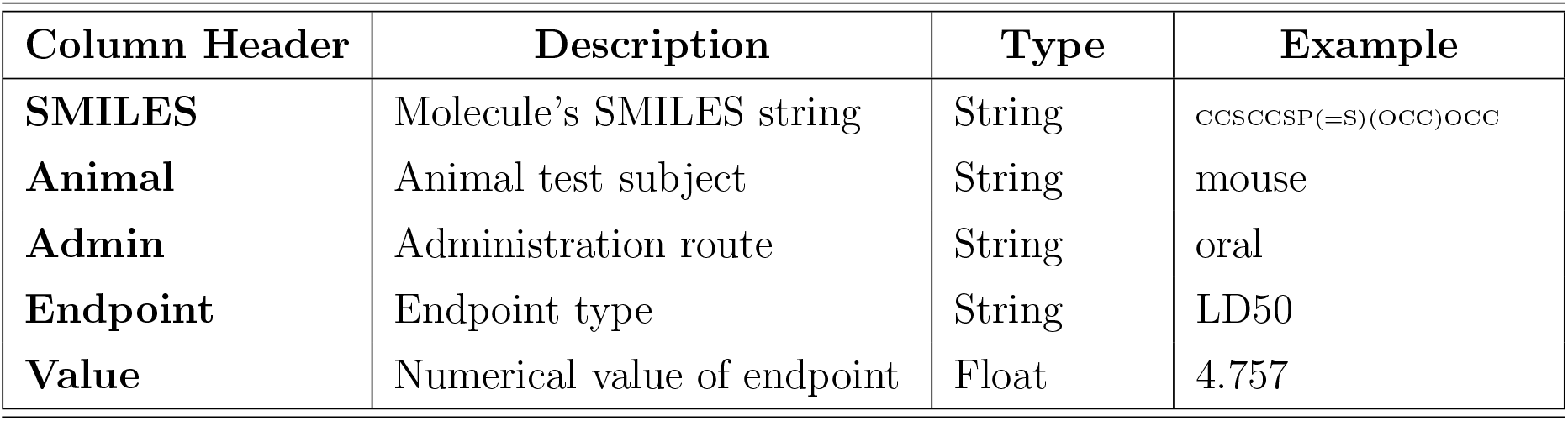
Description of the Canary Dataset Headers

| Column Header | Description | Type | Example |
| --- | --- | --- | --- |
| **SMILES** | Molecule’s SMILES string | String | CCSCCSP(=S)(OCC)OCC |
| **Animal** | Animal test subject | String | mouse |
| **Admin** | Administration route | String | oral |
| **Endpoint** | Endpoint type | String | LD50 |
| **Value** | Numerical value of endpoint | Float | 4.757 |

#### 2.1.2 Molecular Classes of Interest

The Canary dataset contains several molecular classes on which we validated our methods. In particular we consider the “OP” structure-defined class. This class consists of organophos-phate stuctures, many of which are known to target the acetylcholinesterase (AChE) receptor (*n* = 180). We also consider antagonists of the androgen receptor (AR, *n* = 366) and estrogen-receptor-*β* (ER*β*) signaling pathways (*n* = 327) and refer to these target-defined molecular classes as “AR” and “ER*β*”, respectively. AR and ER*β* antagonists were identified by the “AR BLA Antagonist ch2” and “ERb BLA Antagonist ch2” binary endpoints in the Canary dataset. Oral mouse and rat LD_50_ values for identified AR and ER*β* inhibitors were extracted from the Canary dataset for model training.

For comparison, the toxicity distributions for each molecular class of interest are shown in Figure 2-a. We observe that the OP class is overall more toxic than the other classes. Additionally, molecules from these three classes form a distinct cluster relative to the entire Canary dataset (Figure 2-b). The 2-D clustering was obtained by vectorizing the molecular SMILES with embeddings from the baseline model (described in the next section), and then applying Pairwise Controlled Manifold Approximation (PaCMAP)^21^ for dimensionality reduction.

**Figure 2:**
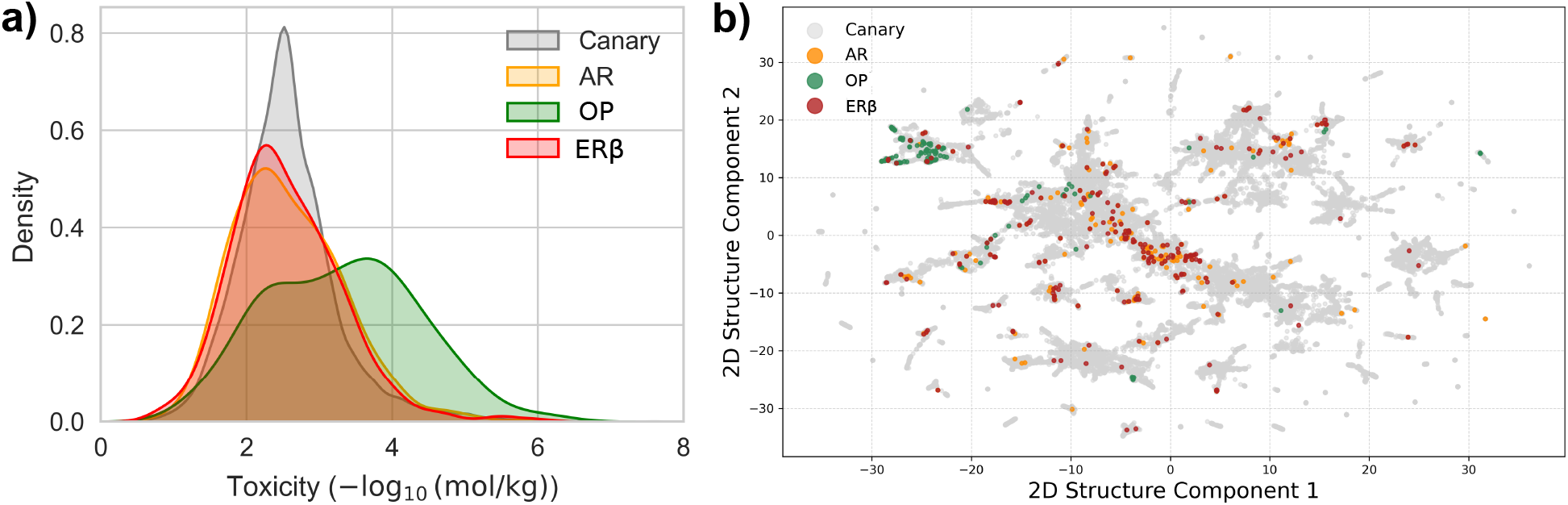
Canary Dataset and Molecular Classes of Interest. (**a**) The oral LD_50_ distribution for the Canary dataset (*n ≈* 28, 000) and for the molecular classes of interest: AR (*n* = 366), ER*β* (*n* = 327), and OP (*n* = 180). These datasets consist of only mouse and rat data. (**b**) 2D visualization of chemical structure for the datasets.

### 2.2 Training Deep Learning Models

The emergence of deep learning ML models in the 2010s revolutionized molecular property prediction, shifting the paradigm from fixed molecular fingerprints to learned fingerprints. Perhaps the most popular and commonly used tool based on learned fingerprints is Chemprop.^17^ The model architecture comprises a directed-graph Message-Passing Neural Network (MPNN) that converts a SMILES string into a fixed-dimensional molecular embedding, followed by a Feedforward Neural Network (FNN) whose final linear layer regresses the property of interest directly from this embedding. The weights of both neural networks are simultaneously varied during training to perform a given classification or regression task. Here, the curated toxicity training set was used to train a Chemprop model using the Chemprop 2.0 package. The overall training pipeline is shown in Figure 3. The overall training and testing details are discussed bellow.

**Figure 3:**
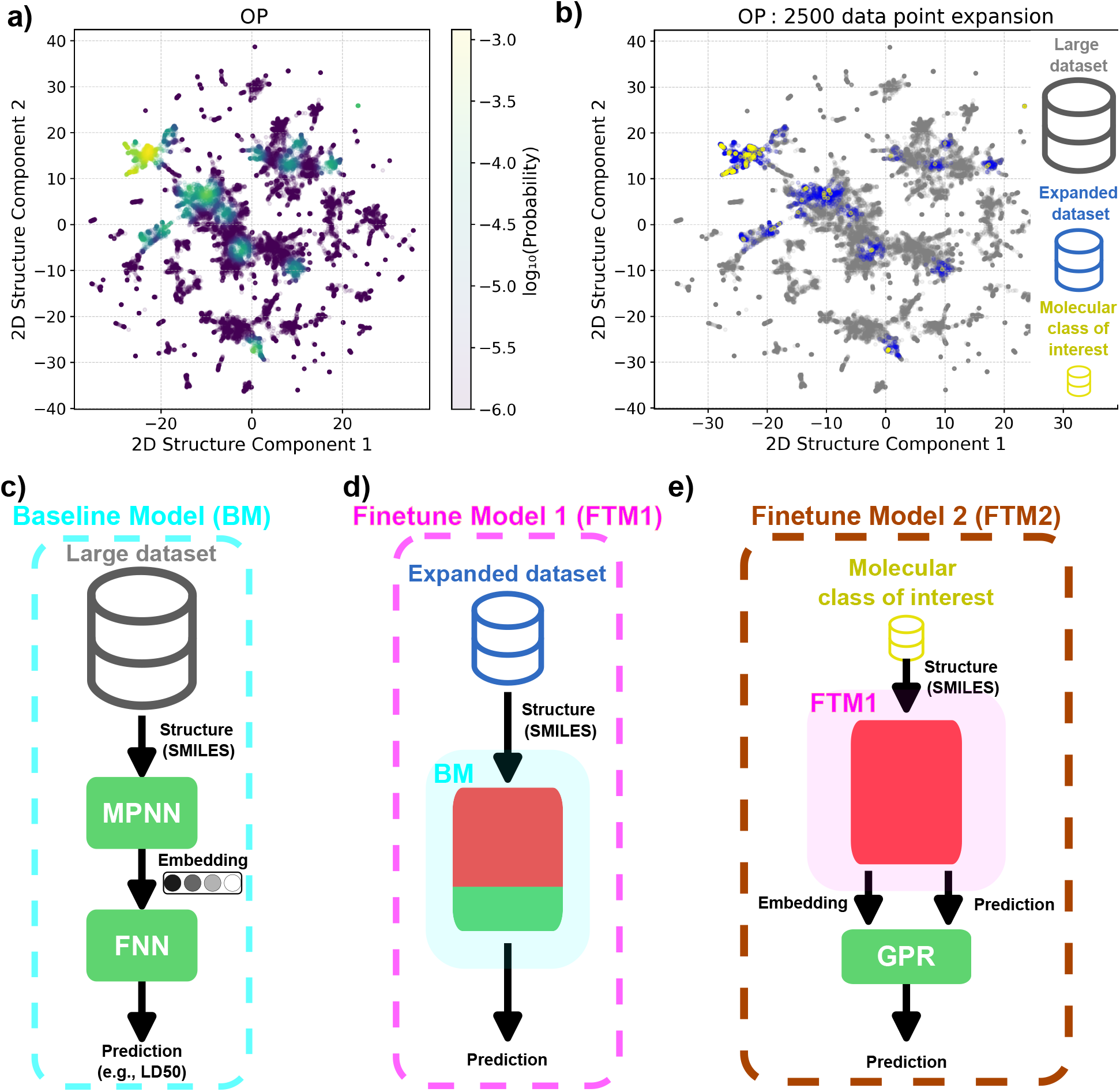
Training and Fine-Tuning Deep Learning Models with a Probabilistic Regressor Head. (**a**) Probability distribution over chemical structure space indicating whether a molecule is a structural analog of compounds in the OP dataset. (**b**) Sampling 2,500 molecules from (a) to define: molecular class of interest (OP), expanded dataset (random samples excluding the class of interest), and large dataset (Canary dataset). (**c**) Baseline model (BM): MPNN generates embeddings from SMILES; FNN regresses target endpoint. Trained on the large dataset. (**d**) Fine-tuned model 1 (FTM1): BM fine-tuned on the expanded dataset with MPNN and some FNN layers frozen. (**e**) Fine-tuned model 2 (FTM2): GPR trained on embeddings and FTM1 predictions from the user-defined molecular class of interest dataset. **Color code**: Red = frozen weights; Green = trainable weights.

#### 2.2.1 Data preprocessing and splitting

**Label scaling.** Toxicity values (the regression target) were z-score normalized:

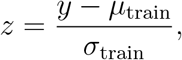

where *µ*_train_ and *σ*_train_ are the mean and standard deviation computed only on the training split. This scaling was applied to all regression learners (RF, XGBoost, SVR, FFNN, etc.). Chemprop performs internal label scaling, so no external normalization was used for that model.

**Data splits.** For each molecular class (structure- or target-defined) we randomly reserved 33% of the compounds as a *held-out test set*, never seen during training or hyper-parameter optimization. The remaining 67% were divided into training and validation subsets using the standard 75%*/*25% split.

#### 2.2.2 Dataset Expansion

We present a sampling strategy that extracts compounds from a large repository (the Canary dataset) based on a user-defined molecular class of interest (e.g., OP). The method identifies and selects molecules whose chemical structures are similar to those that belong to the target class. As described in the following section, the resulting expanded set will be employed for model fine-tuning. The set of sampled molecules is denoted as the expanded dataset. The expanded dataset is constructed by adding structural analogs of the user-defined class molecules; structural similarity is used solely for expansion and does not imply shared target activity. In brief, molecular structures are vectorized using the embeddings from the baseline model (described in the next section), which are then reduced down to two dimensions with the PaCMAP framework. ^21^ Thereafter, the probability distribution of the the user-defined molecular class of interest in chemical structure space is estimated using a Gaussian kernel density estimate on the 2D representations (Figure 3-a). Finally, all molecules in the Canary dataset are assigned a probability label using the model and for a given target size of the expanded dataset, random sampling is performed obeying this distribution (Figure 3-b).

#### 2.2.3 Baseline Model and Optimization

A baseline model is trained using all of the training data in the large dataset (Figure 3-c). The model’s hyperparameters were optimized using this dataset and remained fixed throughout all fine-tuning stages. A discussion of the optimized “basic” hyperparameters, along with the complete model parameters, is provided in SI Section *Optimal Chemprop Architecture*.

The baseline consists of an ensemble of five independently trained replicates; each replicate randomly partitions the training data to create its own validation set. The final prediction for any molecule is obtained by averaging the outputs of the five models. Molecular embeddings used in downstream analyses are extracted from the MPNN’s final hidden layer of the baseline model.

#### 2.2.4 Finetuned Model 1 (FTM1)

A finetuning step is performed on each of the replicates of the baseline model to provide specificity to the the user-defined molecular class of interest (e.g., OP) by leveraging the expanded dataset as the training data. We allowed the size of the expanded dataset to vary between 1,000, 2,500, 5,000, and 10,000 molecules in addition to the class of interest. All the layers of the MPNN along with the first *n* layers (tunable) of the FNN are locked, while the remaining layers are finetuned on the expanded dataset (Figure 3-d).

#### 2.2.5 Finetuned Model 2 (FTM2)

A final fine-tuning step is performed solely on the class of interest to add specificity, while also leveraging the desired properties of a GPR model (Figure 3-e). More precisely, the input to the GPR model is the molecular embedding of the FTM1 (the output of the MPNN) from five replicated models, which are combined by concatenating their outputs, and then subjected to dimension reduction using partial least squares regression (PLSR) with eight components, where the PLSR model is trained on the large dataset (Figure 3-c). The GPR model is trained to predict the error between the true toxicity label and the predictions of the FTM1, along with the variance values of the toxicity predictions from the five replicates of the FTM1 output. This finetuned model adjusts predictions for molecules structurally similar to those belonging to the class of interest while retaining the predictions of the FTM1 for molecules structurally dissimilar to the class of interest. This fine-tuning step is a form of boosting, where the GPR is trained to correct the FTM1’s errors on the class of interest.

### 2.3 Benchmarking Performance

In this study we employed two complementary benchmarking approaches and evaluated them with several performance metrics. First we describe the benchmarks; second we explain how the metrics were calculated.

**Benchmark Workflow with EPA TEST.** To assess the predictive power of our class-specific deep-learning model we compared it with TEST, a model developed by the EPA which is considered a gold-standard QSAR platform.^20^ TEST receives a SMILES string and a chosen toxicity endpoint as inputs and returns regression predictions from a suite of models. For the present work we used the tool’s oral rat LD_50_ endpoint and extracted the prediction from the “consensus” model, which averages the outputs of all individual models. This consensus approach has consistently delivered the best overall performance across diverse chemical datasets.

**Benchmark Workflow with Fixed Fingerprints** For a quick comparison with end-to-end deep learning we used a traditional pipeline that relies on a fixed molecular fingerprint.

1. **Fingerprint Generation:** convert SMILES to an RDKit topological fingerprint (the best performer in our preliminary tests).

## 2. Dimensionality Reduction (Optional): apply Partial Least Squares Regression (PLS) to the fingerprint matrix

1. 3. **Regression:** train five regressors on the (reduced) vectors: Feedforward Neural Network (FNN), Gaussian-Process Regression (GPR), Gradient-Boosted Regression (GBR), Linear Regression (LR) and Support Vector Regression (SVR).

All hyper-parameters (fingerprint size, max-path length, number of PLS components, and regressor-specific settings) were tuned by a grid search, optimizing *R*^2^ with 5-fold cross-validation on the class-specific data. For the full methodological details see SI Section *Benchmark Model Optimization*.

**Performance Metrics.** Model performance was quantified with the *R*^2^ score on a held-out test set for each user-defined molecular class. To facilitate direct comparison with the EPA benchmark we normalized every model’s *R*^2^ by the *R*^2^ obtained from the EPA TEST consensus prediction for the same test set:

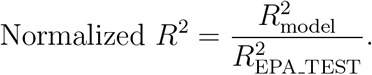

A normalized score greater than 1 indicates performance superior to the EPA benchmark, whereas a value less than 1 denotes inferior performance.

## 3 Results

In this section, we demonstrate that by combining deep learning with a GPR model (FTM2), we can construct class-specific toxicity predictors for structure-defined or target-defined molecular classes that outperform the current gold standard across multiple classes.

### 3.1 Benchmark Model Results

The results of optimizing the Fixed Fingerprint (FP) together with several machine-learning (ML) models are shown in Figure 7 in the Appendix. The performances of the best fixed FP-ML model and of the gold-standard EPA TEST model on the three classes of interest is presented in Table 2. Additionally, we provide the performance of a Chemprop deep-learning model whose architecture was optimized solely for each class-specific dataset. As expected, these Chemprop models exhibit the worst performance when trained on these small data sets (*<* 1000 molecules)^17^ . Notice that *R*^2^ *<* 0 occurs when the residual sum of squares is larger than the total sum of squares, i.e., the model’s error exceeds the data variance. In general, the EPA TEST outperforms the fixed-FP ML benchmark.

**Table 2:** EPA TEST Outperforms Other Benchmark Models on Normalized R. ^2^ **Metric.** R^2^ on the test set normalized to the R^2^ of the EPA TEST benchmark for both the best fixed FP model and a Chemprop model. Results are reported for all three molecular classes of interest (Section 2.1.2). All models are trained and tested solely on the corresponding molecular class of interest

| *Dataset* | *EPA TEST* | *Fixed FP + ML*<br>*Regressor (best)* | *Chemprop*<br>*model* |
| --- | --- | --- | --- |
| *OP* | 1.00 ($R^2 = 0.63$) | 0.70 | 0.63 |
| *AR* | 1.00 ($R^2 = 0.41$) | 0.34 | *Below zero* |
| *ERβ* | 1.00 ($R^2 = 0.43$) | 0.53 | *Below zero* |

**Table 3:** Direct Performance of BM Fine-Tuned with the FTM2 Strategy Relative to the BM. R^2^ on the test set of the BM after the fine-tuning strategy used in FTM2 is applied to it directly normalized to the R^2^ of the BM. Results are reported for all three molecular classes of interest.

| ***Dataset*** | *$R^2_{\text{BM} \rightarrow \text{FTM2}} / R^2_{\text{BM}}$* |
| --- | --- |
| *OP* | 1.021 |
| *AR* | 1.056 |
| *ERβ* | 1.021 |

#### 3.1.1 Finetuned Deep Learning Models

The following provides the results of training and finetunning deep learning models and compares the performance to the best performing EPA TEST benchmark.

**Step 1 - Training the Baseline Model (BM)**: The BM is a Chemprop neural network that was trained exclusively on the training portion of the large dataset (Figure 3). In contrast, the Chemprop model used to generate the benchmark results in Table 2 was trained only on the class-specific training data (i.e., the subset of molecules belonging to the class of interest).

The architecture of the BM (Figure 3-c) was optimized to minimize the cross-validation loss on the entire dataset using the RayTune hyper-parameter tuning module provided with Chemprop. The hyper-parameter values that yielded the best performance are listed in Table 5 of the Supporting Information.

When evaluated on the independent EPA test set, the BM achieves performance that is comparable to the EPA benchmark across all classes (see Table 4).

**Table 4:**
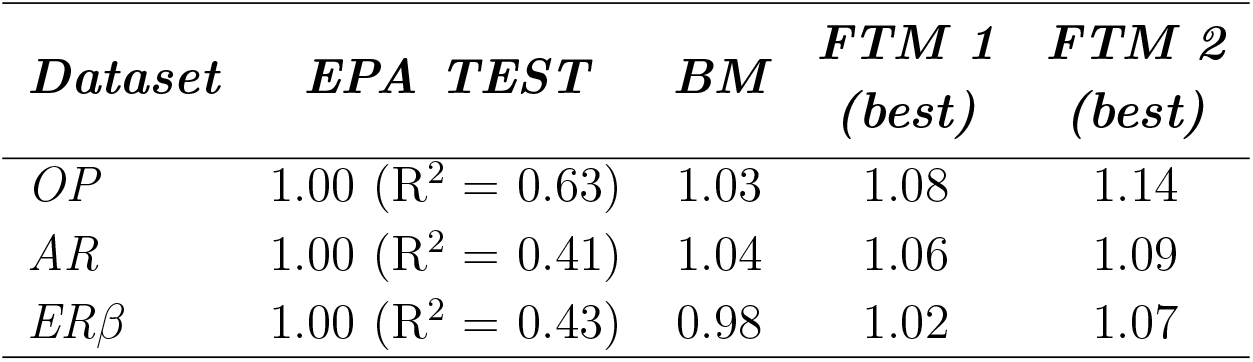
Finetuned Models Perform Better than EPA TEST Gold Standard. R^2^ values on the test set, normalized to the R^2^ of the EPA TEST model, are reported for the BM and the best-performing FTM1 and FTM2 models, with results shown separately for each of the three molecular classes of interest.

| *Dataset* | *EPA TEST* | *BM* | *FTM 1*<br>*(best)* | *FTM 2*<br>*(best)* |
| --- | --- | --- | --- | --- |
| *OP* | 1.00 ($R^2 = 0.63$) | 1.03 | 1.08 | 1.14 |
| *AR* | 1.00 ($R^2 = 0.41$) | 1.04 | 1.06 | 1.09 |
| *ERβ* | 1.00 ($R^2 = 0.43$) | 0.98 | 1.02 | 1.07 |

**Step 2 - Training the Finetuned Model 1 (FTM1)** : The BM was fine-tuned by freezing the layers of the MPNN and allowing some of the weights from the final layers of the FNN to vary with training (FTM1 in Figure 3-d). The goodness of fit on the test sets from fine-tuning on different expansion dataset sizes and by freezing different layers on the FNN are shown in the top diagonals of Figure 4-a,b,c. We observe that the best performance is achieved when all layers of the FNN are frozen except for the final layer. Overall there was not a strong dependence on performance and expansion size.

**Figure 4:**
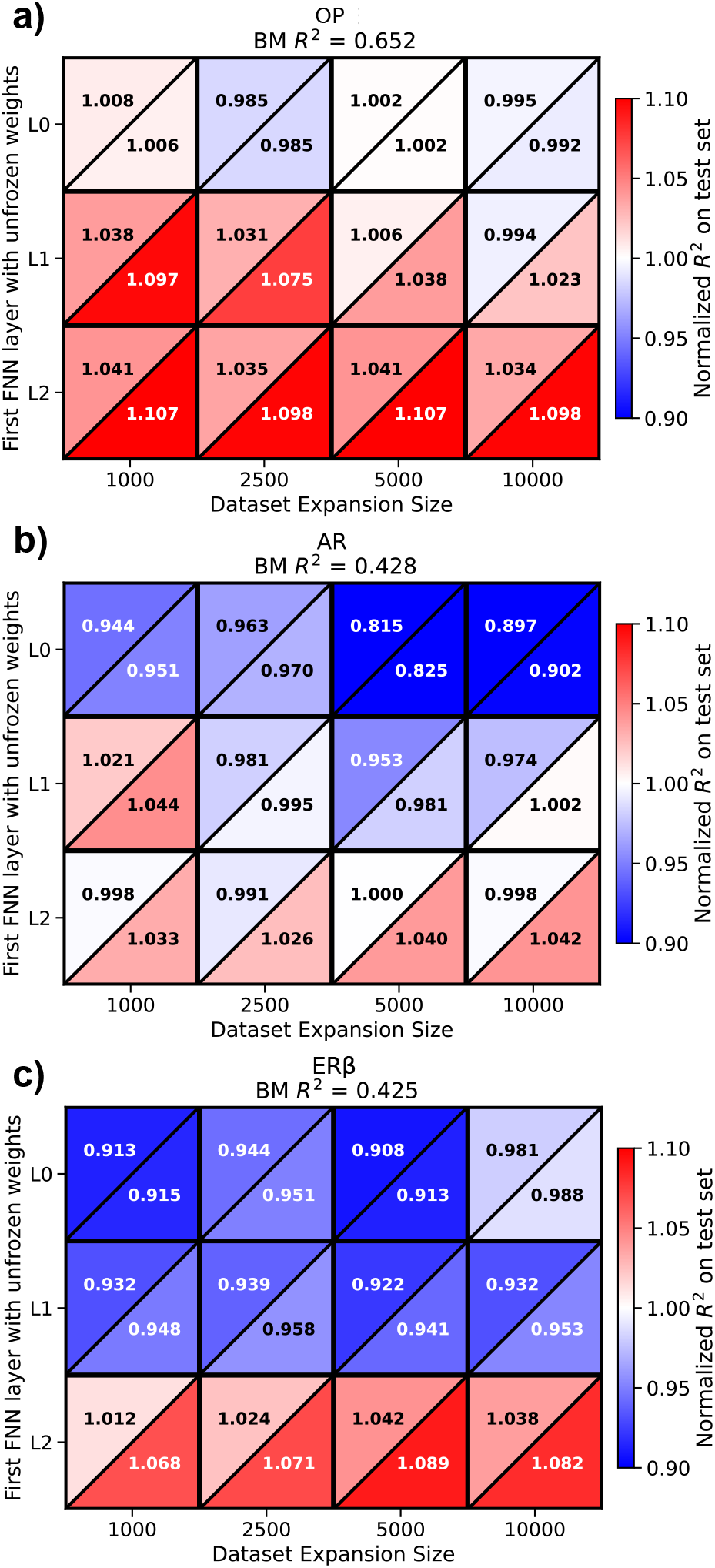
**Fine-Tuning Improves Model Performance Relative to BM**. (**Panels a-c**) Top diagonals show the R² on the test set for FTM1 normalized by the R^2^ of the BM as a function of the number of FNN layers frozen cumulatively from the input layer up to the layer indicated on the y-axis, and the dataset expansion size used for training (x-axis), for the molecular classes: (a) OP, (b) AR, and (c) ER*β*. The lower diagonal shows the R^2^ on the test set (normalized by the R^2^ of the BM) for the model in the top diagonal after the finetuning strategy of FTM2 has been applied to it. The first line of the title denotes the structure-defined or target-defined molecular class, the second line shows the raw R² of the BM.

**Step 3 – Training the Finetuned Model 2 (FTM2).** The second fine-tuning step uses the GPR model to correct the errors made by FTM1 (or by the baseline model, BM) on the molecular class of interest. In this way the overall predictive performance is improved through a boosting strategy (FTM2, see Figure 3-e). The goodness-of-fit for each model after fine-tuning FTM1 is shown in the lower-diagonal panels of Figure 3-a–c. For virtually all models, performance increases after this second fine-tuning stage. We also investigated whether the fine-tuning strategy used in FTM2 could be applied directly to the BM (Table 3), thereby bypassing the intermediate FTM1 step. This results indicate that, except for the AR class (R^2^ of 1.056 vs 1.042 relative to the BM model), the intermediate fine-tuning step (FTM1) is beneficial.

The overall results from all the best trained models, and the benchmarks, are shown in Table 4. Overall, these results suggest that our modeling framework produces class-specific models that outperform the EPA gold-standard benchmark by an average 10% in goodness-of-fit on the held-out test set across the three classes of interest (14% OP, 9% AR, and 7% ER*β*). The GPR head fine-tuned in the second stage boosts specificity for the user-defined molecular class by learning to correct the deep learning model’s errors within that class, while preserving its original predictions elsewhere.

## 4 Conclusions

The recent surge in generative-AI platforms has made it possible to create millions of chemically diverse small-molecule candidates in a matter of hours, dramatically accelerating lead-optimization in drug discovery, agro-chemical design, and related high-impact fields.^1–5^ As these designs drift farther from the scaffolds that underlie existing toxicity models, the predictive reliability of conventional QSAR tools deteriorates, especially when only a few hundred experimentally measured toxicity labels are available.^9–12^

To bridge this gap we introduced a three-stage transfer-learning framework: (i) a graph neural-network baseline is pre-trained on a large public toxicity repository to capture universal chemical knowledge; (ii) the model is fine-tuned on a modest, class-specific dataset (less than 250 molecules per class) using structurally similar compounds to focus the latent space; and (iii) a Gaussian-process regression head is added and jointly fine-tuned to provide probabilistic outputs (Figure 3). Applied to organophosphates, androgen-receptor antagonists and estrogen-receptor-*β* antagonists, the resulting predictors consistently outperformed the EPA’s gold-standard TEST tool, achieving at least ten-percent increase in goodness-of-fit in addition (Table 4) to providing uncertainty on the predictions.^17,20^ The uncertainties also enable active-learning-driven prioritization of compounds for experimental validation, offering a concrete pathway to iterative, cost-effective screening workflows.^13,18^

Beyond the immediate performance gains, the modular nature of the workflow makes it straightforward to incorporate alternative uncertainty-quantification strategies such as evidential deep learning,^22^ deep ensembles^23^ or newer Bayesian methods,^24^ as well as more sophisticated acquisition functions for active learning.^9,13,18^ Enriching the input representation with three-dimensional conformations, docking-derived interaction fingerprints or quantum-chemical descriptors could further boost predictive fidelity for mechanistically complex endpoints.^25,26^ Likewise, replacing the GNN baseline with a pretrained molecular transformer may improve transferability across scaffold-novel classes.^27^ Finally, extending the approach to multi-task or hierarchical models that jointly predict multiple toxicological endpoints promises to reduce the per-task data burden while leveraging shared mechanistic information.^23,28^

In sum, by uniting large-scale pre-training, targeted fine-tuning, and principled probabilistic calibration, we have demonstrated that high-quality, class-specific toxicity predictors can be generated on demand from small, noisy datasets. This capability is poised to become an essential component of data-driven chemical-safety assessments as generative AI continues to expand the frontier of molecular design.

## 5 Data Availability

The data supporting the findings of this study are available from the corresponding author upon reasonable request.

## 6 Acknowledgements

The Authors would like to thank Katherine Schultz, Abdullah Shouaib, Mark Maupin and the rest of the Canary Team at Pacific Northwest National Laboratory (PNNL) for their helpful discussions and the sharing of their Canary dataset which was used in this work. The CANARY work was supported by DHS S&T. The work documented in this publication was funded by the Defense Threat Reduction Agency under Contract HQ003419D0006.

## Supporting Information Available

In this SI we briefly describe (i) the Bayesian-optimization workflow used to identify low-toxicity molecules, (ii) the benchmark-model optimization study that evaluated several classical regressors on fixed-fingerprint features, and (iii) the Chemprop architecture that was selected as the baseline deep-learning model for the three molecular classes of interest.

### 6.1 Bayesian Optimization for Molecular Screening

#### 6.1.1 Motivation and Method

The discovery of low-toxicity compounds from a large chemical library is a classic high-throughput screening problem: testing every molecule experimentally is prohibitively expensive, while random selection of candidates is often inefficient. Bayesian Optimization (BO) offers a principled way to prioritize molecules that are most likely to be non-toxic by iteratively updating a surrogate model and selecting the next batch of compounds with an acquisition function. In this study we implement BO with the Expected Improvement (EI) acquisition function and compare it with pure random sampling. In particular we focus on the OP structure-defined class (*≈* 180 molecules). Of these molecules 60 are randomly held-out for testing and the reminding 120 are used for training.

**Data Preparation.** SMILES strings from the OP dataset are converted to fixed-length fingerprints (RDKit fingerprint with maxpath=3). The Canary dataset with the OP class removed was used to learn a z-score scalar and dimension reduction (partial-least-squares regression with 150 components) to transform the fingerprint vectors. Additionally, the toxicity labels were also z-scored using this dataset.

**Surrogate Model.** A Gaussian Process (GP) with a constant kernel multiplied by an RBF kernel is trained on the currently labelled data. The GP provides a mean prediction *µ*(**x**) and a standard deviation *σ*(**x**) for any candidate **x**.

**Acquisition Function.** For each candidate the Expected Improvement (EI) is computed as

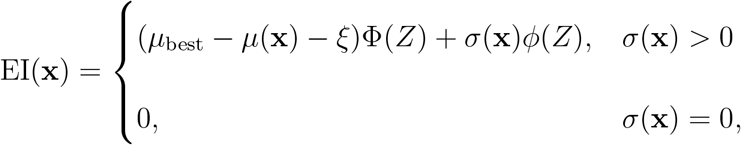

where *Z* = (*µ*_best_ *−µ*(**x**)*−ξ*)*/σ*(**x**), Φ and *ϕ* are the Cumulative Distribution Function (CDF) and Probability Density Function (PDF) of the standard normal distribution, respectively, and *ξ* is a small exploration parameter (set to 10*^−^*^6^ in the experiments).

**Active-Learning Loop.** At each iteration a batch of *b* molecules (here *b* = 3) is selected:

- **Random Strategy:** *b* molecules are drawn uniformly from the remaining pool.
- **EI Strategy:** the *b* molecules with the highest EI scores are chosen.

The selected molecules are added to the training set, the GP is refitted, and performance metrics are recorded on an independent test set. The loop continues until the pool is exhausted or a predefined number of iterations is reached. The whole procedure is repeated *N*_trials_ = 1000 times to obtain reliable statistics.

#### 6.1.2 Results

The BO approach substantially outperforms random sampling in two respects: it discovers low-toxicity molecules more quickly, and it predicts the toxicity of those low-toxicity molecules more accurately.

- **Model Performance on the Full Test Set** (Figure 5a) shows a faster increase in *R*^2^ for the EI strategy; the mean *R*^2^ reaches *≈* 0.20 after screening *∼* 60 molecules, whereas random sampling reaches the same plateau *R*^2^ but requires all 120 molecules.
- **Mean Toxicity of the Lowest 10% of Screened Molecules** (Figure 5b) declines more rapidly under EI, indicating that the EI policy concentrates on low-toxicity regions of chemical space.
- **Best (Lowest) Toxicity Discovered** (Figure 5c) is consistently lower for EI than for random sampling; on average the lowest-toxicity molecule is found after screening roughly 60 molecules with EI, whereas random sampling requires about 120 molecules.
- **Prediction Error on Low-Toxicity and High-Toxicity Subsets** (Figure 5d and 5e) shows that EI achieves a considerably smaller Mean-Squared Error (MSE) for the low-toxicity 10^th^ percentile than random sampling average MSE with fewer molecules screened. For the high-toxicity 90^th^ percentile the random sampling method yields a lower MSE than the EI.

**Figure 5:**
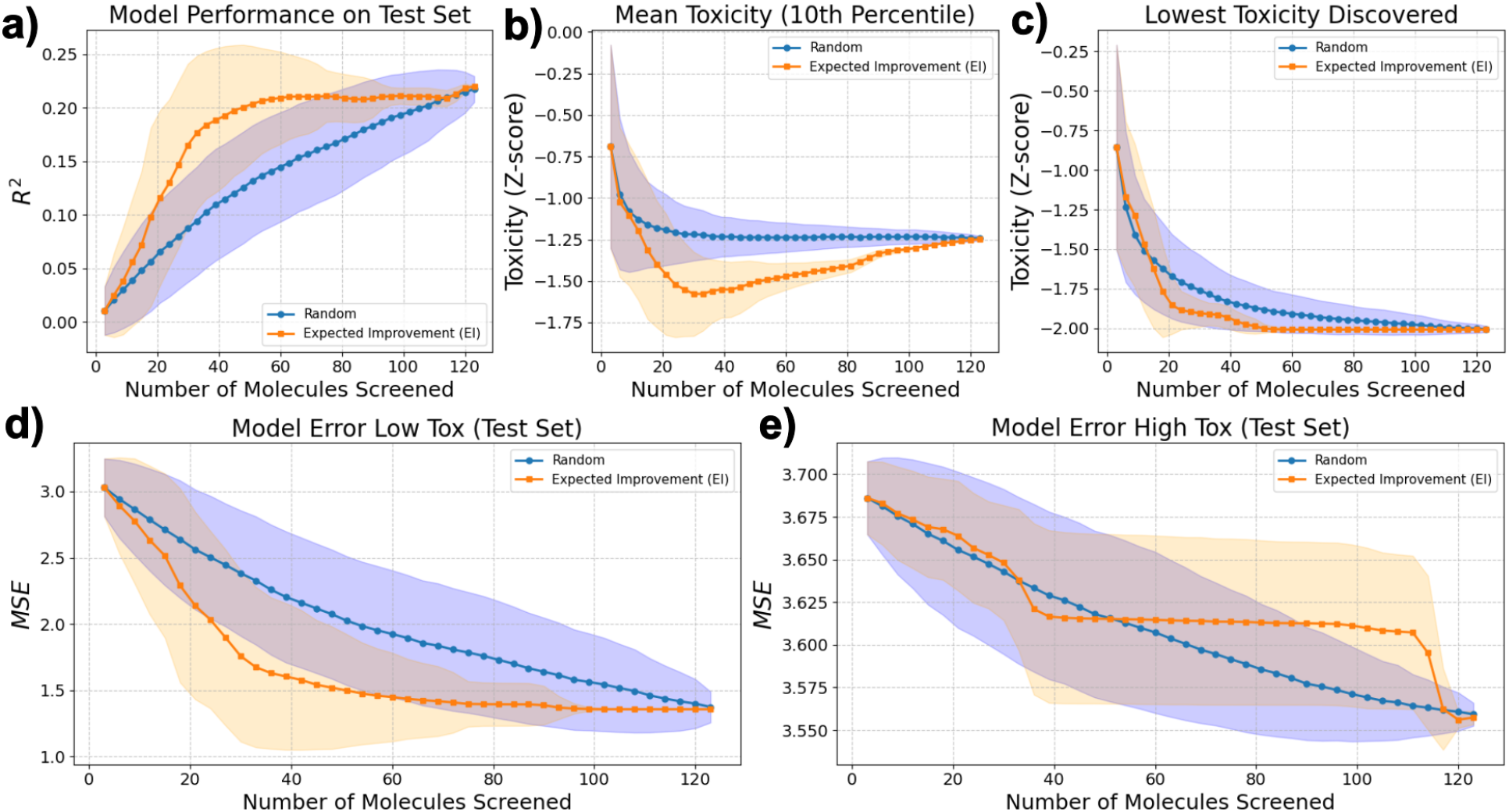
The Model Rapidly Identifies Low-Toxicity Molecules from a Pool of Unknown Compounds. (**a**) Goodness-of-fit of the model on a held-out test set as the number of screened molecules increases. (**b**) Mean toxicity of the molecules that fall within the lowest 10th percentile of the screened set. (**c**) Toxicity value of the single least-toxic molecule discovered. (**d**) Mean-Squared Error (MSE) on a held-out test set for the lowest-toxicity 10% of molecules (*≤* 10th percentile). Orange curves correspond to Bayesian-optimization sampling, while the blue curve represents random sampling. (**e**) MSE on a held-out test set for the highest-toxicity 10% of molecules (*≥* 90th percentile). Shaded regions indicate one standard deviation computed over 1,000 repeated experiments.

**Figure 6:**
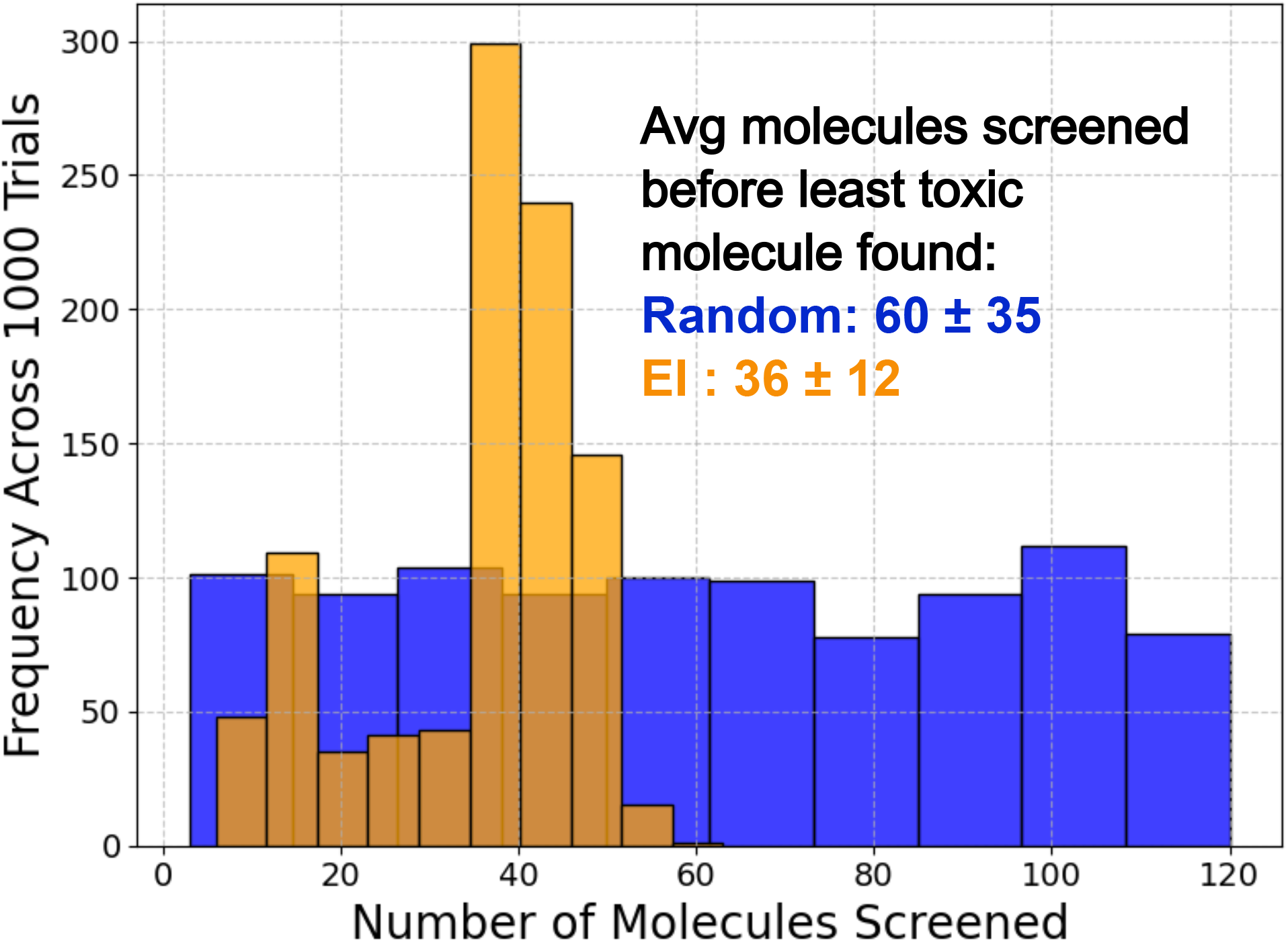
Distribution of How Quickly the Lowest-Toxicity Molecule is Found. The histogram shows the number of molecules screened before the lowest-toxicity compound is identified from a pool of unlabeled molecules. Results are shown for an active-learning strategy (orange) and for random sampling (blue). The distribution is based on 1,000 repeated experiments.

**Figure 7:**
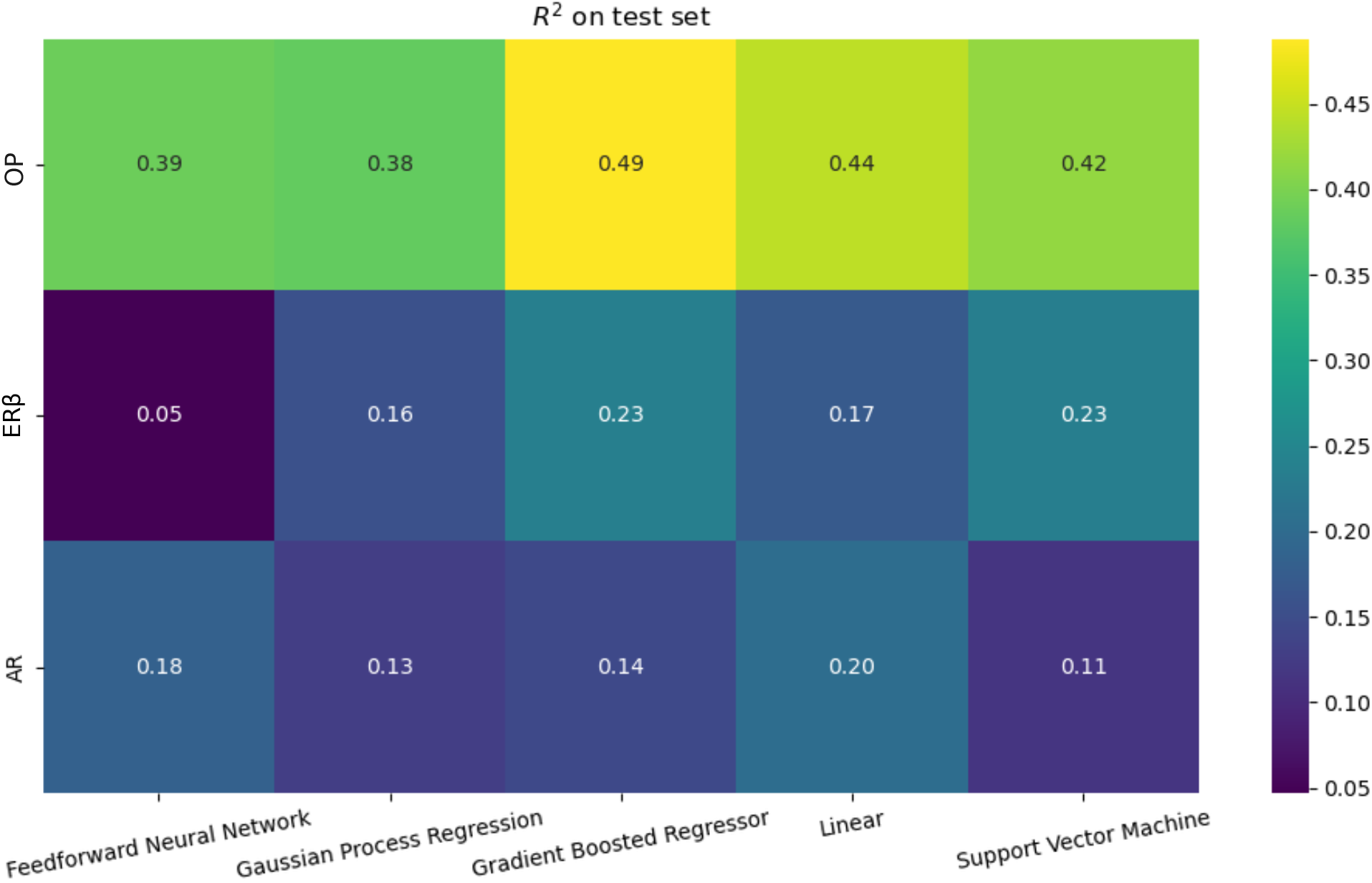
Training of Baseline Fixed Molecular Fingerprint ML Models Across the Molecular Classes of Interest. Test-set R^2^ values obtained with RDKit-generated molecular fingerprints and the following regressors: neural network, Gaussian process regression, boosted-tree ensemble, linear regression, and support-vector regression (Section 2.6.2).

Figure 6 presents the distribution of the number of molecules that must be screened before the globally least toxic compound is found. The EI histogram (orange) is shifted toward smaller numbers; the median number of screened molecules is 36 for EI versus 60 for random sampling. Across 1 000 trials the EI strategy locates the optimal molecule after screening approximately 30% of the pool, while random sampling requires nearly half the entire pool.

These results demonstrate that Bayesian optimization with the Expected Improvement acquisition function provides a rapid and reliable route to identify low-toxicity candidates, dramatically reducing the experimental burden compared with näıve random screening.

### 6.2 Benchmark Model Optimization

#### 6.2.1 Motivation and Method

For each of the three molecular classes (OP, AR, ER*β*) we built a scikit-learn pipeline that consists of

1. **Fingerprinting:** RDKit fixed-length fingerprints (FP Fixed) with two hyper-parameters explored: maxpath = *{*2, 4, 8, 16*}* and fingerprint length fp size = *{*150, 500, 1000, 2000*}*.
2. Scaling: standardization of the fingerprint vectors.
3. **Dimensionality Reduction (Optional):** either no reduction (“passthrough”) or a partial-least-squares (PLS) projection with the number of components drawn from *{*5, 10, 20, 50, 100, 150, all*}*.
4. **Regressor:** one of five widely used models:

- linear regression
- support-vector regression (grid on *C* = *{*0.1, 1, 10, 100*}* and *ɛ* = *{*0.01, 0.1, 0.5, 1*}*)
- multi-layer perceptron (several hidden-layer configurations, *α* = *{*1e*−*4, 1e*−*3, 1e*−*2*}*, learning-rate = *{*constant, adaptive*}*)
- gradient-boosted trees (grid on number of estimators = *{*10, 25, 50*}*, learning-rate = *{*0.01, 0.1, 1*}*, max-depth = *{*3, 5, 10*}*, min-samples-split = *{*2, 5, 10*}*, min-samples-leaf = *{*1, 2, 4*}*)
- Gaussian-process regression with a constant*×*RBF kernel plus white-noise (kernel variance, length-scale and noise level tuned)

For every combination of these settings a 5-fold cross-validation grid-search (scoring = *R*^2^) was performed; the best model for each regressor-class pair was saved for later use.

#### 6.2.2 Results

Figure 7 summarizes the test-set *R*^2^ values obtained after the exhaustive grid-search. For the OP class the best *R*^2^ is *≈* 0.49, for the ER*β* class it is *≈* 0.23, and for the OP class it is *≈* 0.20. Overall, the grid-search provides a benchmark view of the performance ceiling of a fixed-fingerprint approach when combined with several state-of-the-art regressors for each class of interest.

### 6.3 Optimal Chemprop Architecture

The aim of this brief study was to identify a Chemprop configuration that performs well for the Baseline Model (BM) when trained separately on each of the two target-specific (AR, ER*β*) and one structure-specific (OP) molecular classes of interest. To this end we used the Canary data set, excluding the held-out test set, for all three molecular classes.

**Search Strategy.** Chemprop’s built-in hyper-parameter optimization (“hpopt”) was employed. The optimizer is powered by Ray Tune. For every data set we performed

- 5 independent replicates (“–num-replicates 5”) – each replicate uses a fresh random 80/10/10 train/validation/test split;
- 500 trials per replicate (“–raytune-num-samples 500”);
- early-stopping after 45 epochs with a reduction factor of 2.

The search space was limited to the six parameters that most strongly influence a message-passing neural network and that are included in the Chemprop key work *basic*.

1. depth - number of message-passing steps.
2. ffn num layers - number of fully-connected layers after message passing.
3. dropout - dropout probability in the MPNN and FFN.
4. message hidden dim - hidden dimension of the message vectors.
5. ffn hidden dim - hidden dimension of the feed-forward network.
6. learning-rate schedule (init lr, max lr, final lr, warmup epochs).

All other Chemprop options were left at their defaults. The objective function was validation *R*^2^ (“tracking-metric = val loss”).

**Resulting Configuration.** Across the five replicates the same hyper-parameter set consistently gave the highest validation score for each of the three classes. The configuration is reproduced in Table 5.

**Table 5:**
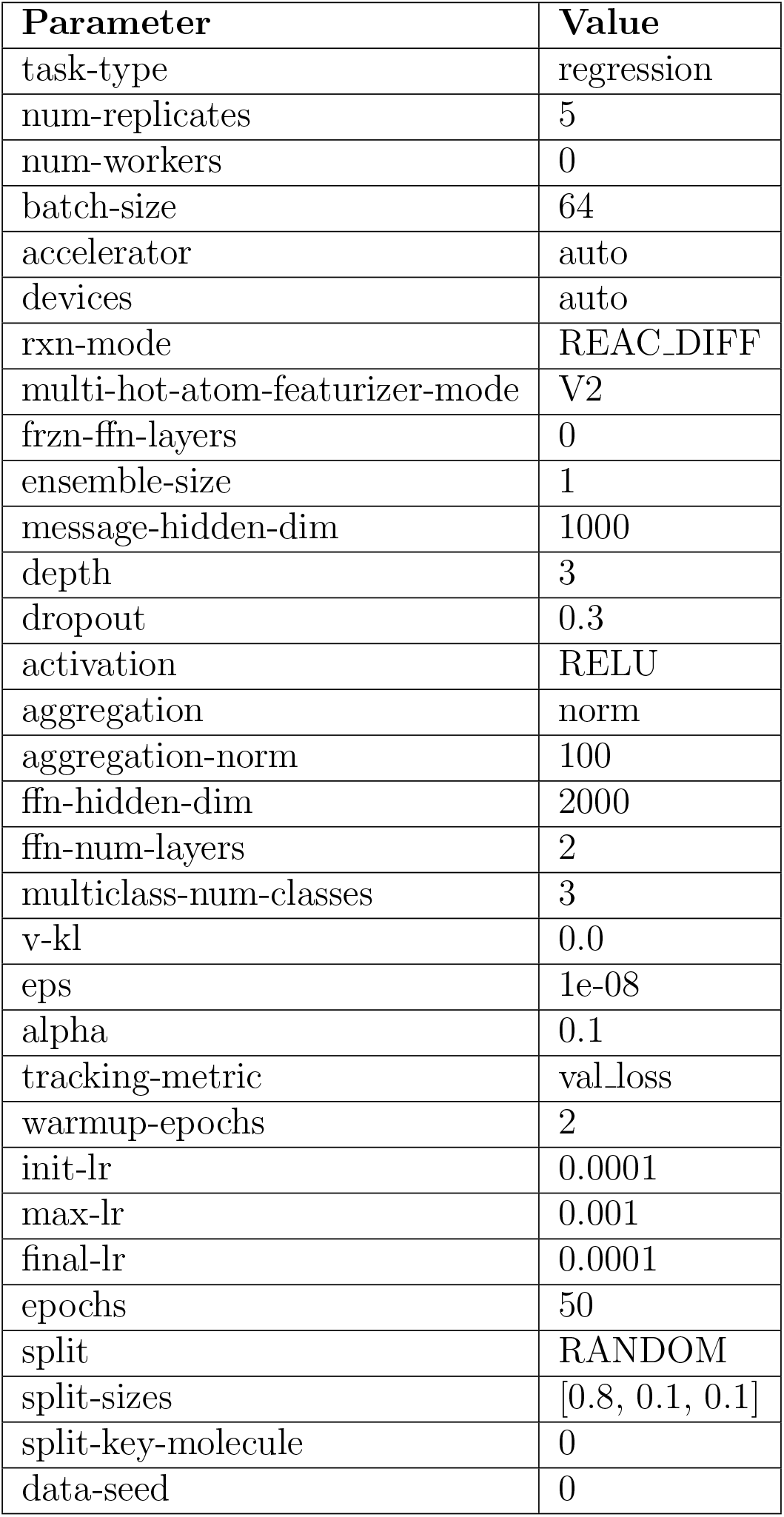
Optimal Chemprop Configuration (identical for the three toxicity data sets).

